# The Pentose Phosphate Pathway constitutes a major metabolic hub in pathogenic *Francisella*

**DOI:** 10.1101/2021.01.15.426780

**Authors:** Héloise Rytter, Anne Jamet, Jason Ziveri, Elodie Ramond, Mathieu Coureuil, Pauline Lagouge-Roussey, Daniel Euphrasie, Fabiola Tros, Nicolas Goudin, Cerina Chhuon, Ivan Nemazanyy, Fabricio Edgar de Moraes, Carlos Labate, Ida Chiara Guerrera, Alain Charbit

**Affiliations:** Université de Paris, 75015 Paris, France; INSERM U1151 - CNRS UMR 8253, Institut Necker-Enfants Malades. Team 7: Pathogénie des Infections Systémiques, 75015 Paris, France; Plateforme Protéome Institut Necker, PPN, Structure Fédérative de Recherche Necker INSERM US24-CNRS UMS 3633; Plateforme Etude du métabolisme, Structure Fédérative de Recherche Necker INSERM US24-CNRS UMS 3633; Pole Bio-analyse d’images, Structure Fédérative de Recherche Necker INSERM US24-CNRS UMS 3633; Laboratório Max Feffer de Genética de Plantas, Departamento de Genética, Escola Superior de Agricultura Luiz de Queiroz, Universidade de São Paulo, Piracicaba, Brazil

**Keywords:** Pentose Phosphate Pathway, metabolic adaptation, intracellular parasitism, *Francisella tularensis*

## Abstract

Metabolic pathways are now considered as intrinsic virulence attributes of pathogenic bacteria and hence represent potential targets for anti-bacterial strategies. Here, we addressed the role of the pentose phosphate pathway (PPP) and its connections with other metabolic pathways in the pathophysiology of *Francisella novicida*. The involvement of the PPP in *Francisella* intracellular life cycle was first demonstrated with the study of PPP inactivation mutants. Indeed, inactivation of *tktA, rpiA* or *rpe* genes, severely impaired intramacrophagic multiplication during the first 24 hours. Time-lapse video microscopy demonstrated that *rpiA* and *rpe* mutants were able to resume late intracellular bacterial multiplication. To get further insight into the links between the PPP and other metabolic networks of the bacterium, we next performed a thorough proteo-metabolomic analysis of these mutants. We show that the PPP constitutes a major bacterial metabolic hub with multiple connections with glycolysis, tricarboxylic acid cycle and other pathways, such as fatty acid degradation and sulfur metabolism. Hence, our study highlights how the PPP is instrumental to *Francisella* pathogenesis and growth in its intracellular niche.

## Introduction

*Francisella tularensis* is the causative agent of the zoonotic disease tularemia (Sjöstedt, 2011). This facultative intracellular bacterial pathogen is able to infect numerous cell types but is thought to replicate and disseminate mainly in macrophages *in vivo* (Santic *et al*, 2006). The four major subspecies (subsp) of *F. tularensis* currently listed are the subsps: *tularensis, holarctica, mediasiatica* and *novicida* (the latter is also called *F. novicida*). These subsps differ in their virulence and geographical origin (McLendon *et al*, 2006) but all cause a fulminant disease in mice that is similar to tularemia in humans (Kingry & Petersen, 2014). Although *F. novicida* is rarely pathogenic in humans, its genome shares a high degree of nucleotide sequence conservation with the human pathogenic subsp *tularensis* and is thus widely used as a model to study highly virulent subspecies.

*Francisella* virulence is tightly linked to its capacity to multiply exclusively in the cytosolic compartment of infected cells, and in particular in macrophages in vivo. Cytosolic pathogens, that notably include *Listeria monocytogenes* and *Shigella flexneri*, often require the utilization of multiple host-derived nutrients (Lobato-Márquez *et al*, 2019; Kentner *et al*, 2014; Keeney *et al*, 2007) and hexoses are generally their preferred carbon and energy sources. The capacity of *Francisella* to multiply in the host cytosol is controlled by multiple regulatory circuits (Ziveri *et al*, 2017), connected to metabolism. In particular, we have shown that gluconeogenesis was essential for *Francisella* intracellular multiplication (Brissac *et al*, 2015; Ziveri *et al*, 2017b), allowing host-derived substrates such as amino acid, pyruvate and glycerol to be used as carbon, nitrogen and energy sources.

The pentose phosphate pathway (PPP) constitutes, with glycolysis, a major pathway for glucose catabolism. However, its contribution to bacterial metabolic adaptation and especially its importance in bacterial pathogenesis, remains largely unexplored. The PPP is composed of two branches, an oxidative and a non-oxidative branch (Ge *et al*, 2020). Glucose flux through the oxidative branch produces NADPH, an essential reductant in anabolic processes. The non-oxidative branch generates the five-carbon sugar Ribose-5P (R-5P) from glucose and can be reversibly converted into glycolytic intermediates *e*.*g*. glyceraldehyde 3P (GA-3P) and Fructose-6P (F-6P). *Francisella*, which lacks the oxidative branch of the PPP, is equipped with a complete non-oxidative branch composed of the four enzymes: *tktA* (*FTN_1333*, encoding a transketolase), *rpiA* (*FTN_1185*, encoding a ribose 5-phosphate isomerase), *rpe* (*FTN_1221*, encoding a ribulosephosphate 3-epimerase) and *tal* (*FTN_0781*, encoding a transaldolase) (**Appendix Fig S1**). These four genes are scattered along the *F. novicida* chromosome and each of them is present in a single copy.

Transketolase, which is central to the non-oxidative branch of the PPP, can produce either R-5P or Sedoheptulose-7-phosphate (S-7P) in response to available metabolite concentrations. Ribulose-5-phosphate (Ru-5P), can be converted either into xylulose-5-phosphate (Xyl-5-P) by Rpe or R-5P by Rpi, respectively. Finally, Tal catalyzes the conversion of S-7P and GA-3P to erythrose-4-phosphate (E-4P) and fructose-6-phosphate (F-6P). Overall, the different precursors synthesized by the non-oxidative branch of the PPP contribute to multiple important functions of the bacterial cell, including biosynthesis of lipopolysaccharide (LPS), aromatic amino acids and nucleic acid precursors.

We performed here the functional analysis of the four PPP mutants (*tkt, rpe, tal* and *rpiA*) and a global network-based approach of their proteo-metabolomic characteristics to highlight pathways related to the PPP and potential biological hubs. The data presented suggest a biphasic bacterial regulation mode between glycolysis and the PPP during intracellular multiplication, and reveal previously unrecognized links between the PPP and other metabolic pathways.

## Results

### Transketolase, a conserved enzyme of the PPP

Transketolase enzymes are ubiquitously expressed in eukaryotes and bacteria. In bacteria, they allow the production of precursors required for the synthesis of nucleotides and certain amino acids as well as for LPS synthesis (**Appendix Fig S1**). Several bacteria express multiple isoforms of transketolases. For example, *Escherichia coli* possesses two genes (*tktA* and *tktB*) and *Salmonella typhimurium*, three genes (*tktA, tktB* and *tktC*), encoding transketolases with different enzymatic properties. *Citrobacter rodentium* genomes encode up to six isoforms of transketolases. In contrast, *Francisella* genomes possess a unique transketolase-encoding gene (here designated *tktA*). The transketolase TktA of *F. novicida* (FTN_1333) is a 663 amino acid long protein that shows 41% to 57% amino acid sequence identity with its orthologs in other pathogenic bacterial species (*e*.*g*. it shares 55.6% and 41% amino acid identity with the transketolases of *Legionella pneumophila* and *Mycobacterium tuberculosis*, respectively).

We have previously shown that *tktA* is the first gene of a highly conserved operon among *F. tularensis* species (Ziveri *et al*, 2017), preceding genes (*gapA, pgK, pyK* and *fba*, respectively) involved in glycolysis/gluconeogenesis (**Fig 1A, B**). The organization of the four first genes of this operon (*tktA-pyK*) is conserved in both *L. pneumophila* (Häuslein *et al*, 2017) and *Coxiella burnetii* species (**Appendix Fig S2** and **Appendix Table S1**). Several other pathogenic bacterial species, such as *Bordetella pertussis* and *Brucella melitensis*, also have *tktA, gapA* and *pgk* genes in the same genetic cluster, and with the same organization, but lack the *pyk* and *fba* genes. Notably, in most *Burkholderia* species (including *B. multivorans, B. pseudomallei, B. thailandensis*, …), the *tktA* and *gapA* genes are adjacent and in the same orientation, suggesting that they belong to the same transcriptional unit, whereas the *pgk, pyk*, and *fba* genes cluster is located in a distinct region of the chromosome (**Appendix Fig S2** and **Appendix Table S1**). These observations prompted us to first evaluated how the expression of the two consecutive genes *tktA* and *gapA* of *F. novicida* would be infleunced by the presence of a glycolytic or a gluconeogenic carbon source. For this, we quantified their transcription by qRT-PCRs in wild-type bacteria grown in chemically defined medium (CDM) supplemented either with glucose (CDM-Glucose) or glycerol (CDM-Glycerol) (**Fig 1C**). Transcription of each gene was roughly similar in the presence of either glucose or glycerol, suggesting that expression of *tktA* and *gapA* is not under the control of these carbon sources. To evaluate a possible growth phase-dependence effect, we also compared expression of *tktA* and *gapA* genes during early (OD_600nm_ of 0.5) or late (OD_600nm_ of 1-1.2) exponential phase of growth. In CDM-Glucose, *gapA* gene expression significantly increased in late exponental phase compared to early exponential phase wheras expression of *tktA* remained unchanged. In CDM-Glycerol, a slight but non-significant increased expression of both genes was observed. Of note, with both carbon sources, expression of *gapA* gene was higher than that of *tktA* (up to 2.5-fold higher in late exponential phase, in CDM-Glucose), suggesting that *gapA* possesses its own promoter.

**Figure 1.**
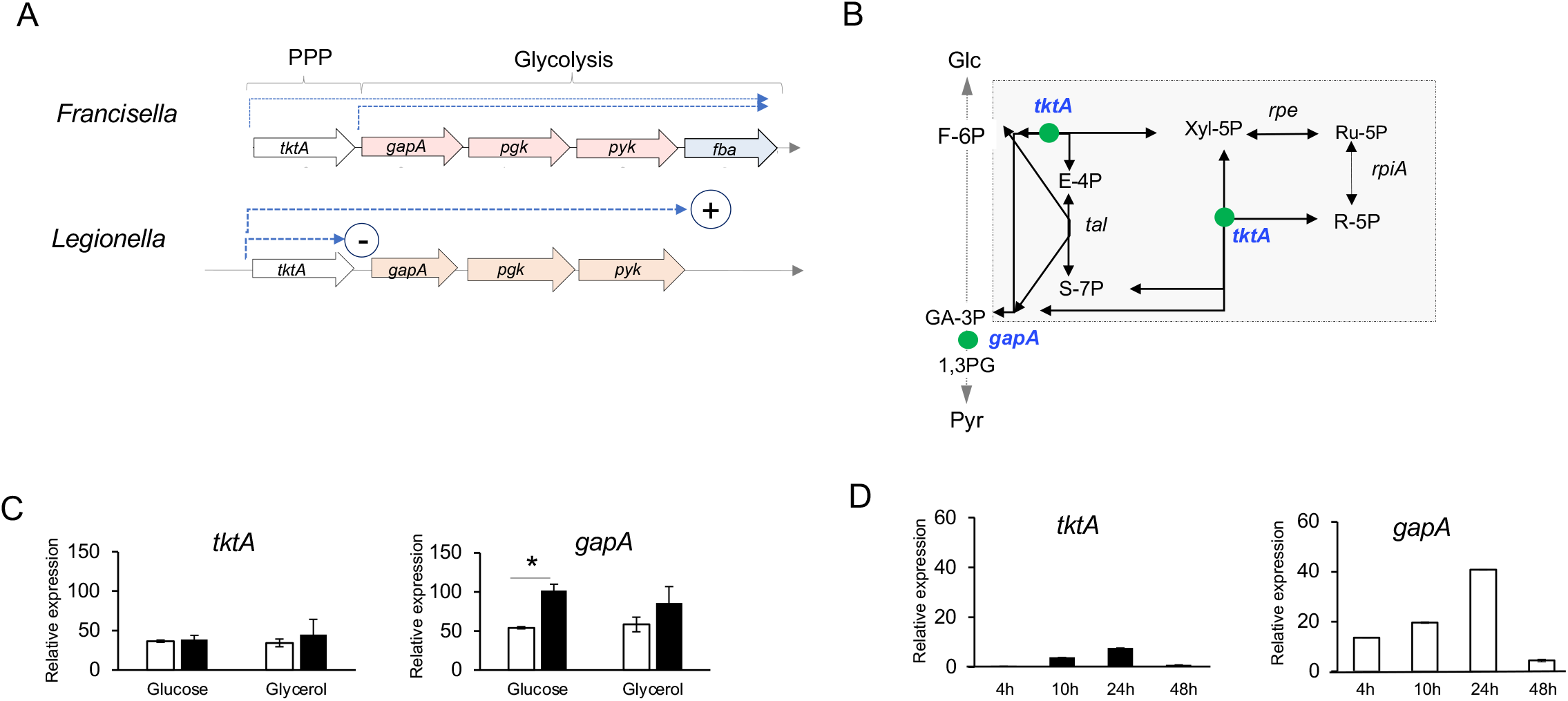
Transcriptional analysis of *tktA* and *gapA* genes. **(A)** Schematic organization of the *tktA* operon in *F. novicida* and *Legionella pneumophila*. The terms PPP and Glycolysis on top, indicate the genes involved in either the PPP or the Glycolytic/gluconeogenic pathways. The last gene of the *Francisella* locus, absent in the *Legionella* locus (*fba*). The dotted blue arrows indicate the predicted transcriptional units. In *Legionella*, in the absence of the CsrA regulatory protein, transcription is interrupted after *tktA* (circled sign -) whereas in the presence of CsrA, transcription resumes till the end of the operon (circled sign +). (B) The enzymatic reactions corresponding to TktA and GapA enzymes are shown as green balls on a schematic depiction of the PPP and glycolytic pathways. **(C)** qRT-PCR analysis of *tktA* and *gapA* genes in WT *F. novicida*, grown in CDM supplemented either with glucose or glycerol, in exponential and stationary phase. **(D)** qRT-PCR analysis of *tktA* and *gapA* genes in WT *F. novicida*, over a 24 hour-period of intracellular growth in J774-1 macrophages.

We next quantified the transcription of *tktA* and *gapA* genes in J774.1 macrophages infected by wild-type *F. novicida*, during a 48 h-period in DMEM-Glucose (**Fig 1D**). In these conditions, expression of both genes progressively increased during the first 24 h (corresponding to the active phase of intracellular bacterial mulutiplication), and dropped at 48 h. Of note, at all time-points tested, *gapA* gene expression was significantly higher than that of *tktA* (approximately 5-fold higher), supporting the notion of an expression of *gapA* from its own promoter.

### Transketolase inactivation impairs bacterial growth and intracellular multiplication

Earlier studies have already shown that transketolase mutants of *Francisella* were defective for growth in macrophages and attenuated for virulence in mice (Tempel *et al*, 2006; Su *et al*, 2007; Asare & Abu Kwaik, 2010; Moule *et al*, 2010). It was also recently reported that inactivation of the *tktA* gene in *F. novicida* drastically reduced the formation of outer membrane vesicles production during the stationary phase of growth in broth, linking the PPP with vesicles biogenesis and nutritional stress (Sampath *et al*, 2018). To extend these findings, we constructed an isogenic deletion mutant of *tktA* in *F. novicida* (Δ*tktA*) by allelic replacement (see Materials and Methods) and a complemented strain (Δ*tktA*-Cp) carrying a plasmid-borne wild-type *tktA* allele. We first evaluated the impact of *tktA* inactivation on bacterial growth in Chemically Defined Medium (CDM) (Chamberlain, 1965), supplemented with various carbohydrates as well as in complex media (Tryptic soy broth and Schaedler K3). Growth of the Δ*tktA* mutant was severely impaired in all the CDMs tested (**Fig 2A**) but wild-type growth was always restored in the complemented strain (Δ*tktA*-Cp), demonstrating the absence of polar effect of the mutation. In complex media such as Tryptic soy broth supplemented with cysteine and glucose (TSB) or Shaedler K3 medium, growth of the Δ*tktA* mutant was restored to almost wild-type levels. Hence, although the PPP seems to be essential for growth in synthetic media, *tktA* is not an essential gene in *F. novicida*.

**Figure 2.**
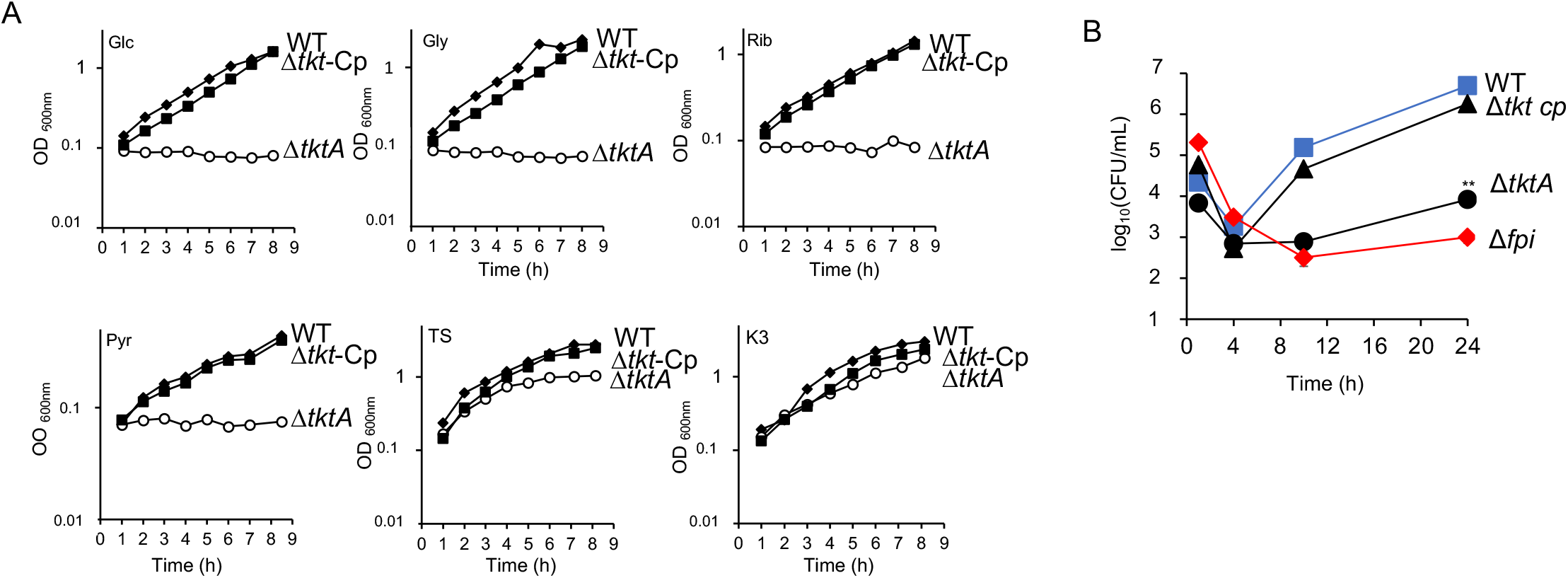
Growth of the Δ*tktA* mutant in liquid culture. (**A**) Bacterial growth was monitored in chemically defined medium (CDM), supplemented with various carbon sources (Glc, glucose; Gly, glycerol; Rib, ribose; Pyr, pyruvate,), each at a final concentration of 25 mM; as well as in tryptic soy broth supplemented with cysteine (0.1% w/v) and glucose (0.4% w/v) (TS); or Shaedler K3 medium (K3). Stationary-phase bacterial cultures of wild-type *F. novicida* (WT), Δ*tktA* mutant and complemented strain (Δ*tktA-Cp*), were diluted to a final OD_600nm_ of 0.1, in 20 mL broth. Every hour, the OD_600nm_ of the culture was measured, during a 9 h-period. (**B**) Intracellular multiplication of the Δ*tktA* and *tktA*-complemented (ΔtktA cp) strains was monitored in J774.1 macrophages over a 24 h-period in DMEM supplemented with Glucose and compared to that in the wild-type *F. novicida* (WT). A Δ*fpi* mutant strain was used as a (A) negative control. ∗∗, P <0.01 (as determined by Student’s t test).

We next evaluated the impact of *tktA* inactivation on bacterial multiplication in the murine macrophage-like cell line J774.1 macrophages (**Fig 2B**) in DMEM-Glucose. In agreement with earlier reports, intracellular multiplication of the Δ*tktA* mutant was totally abrogated during the first 10 h after infection and similar to that of the Δ*fpi* mutant lacking the entire *Francisella* pathogenicity island (Weiss *et al*, 2007). However, a modest increase of bacterial counts (CFUs) was recorded at 24 h, indicating that the Δ*tktA* mutant had started to multiply (an 8-fold increase in CFUs compared to Δ*fpi*). Functional complementation (*i*.*e*. introduction of a plasmid-borne wild-type *tktA* allele into the Δ*tktA* mutant strain) restored wild-type growth.

Altogether, these data showed that the ability mediated by the transketolase TktA to consume the carbon sources available in the cytosol of infected macrophages is essential for intracellular multiplication.

### Contribution of the other PPP enzymes to bacterial growth and intracellular multiplication

We further evaluated the contribution of the PPP to *Francisella* pathogenesis by monitoring the impact of mutations in the three other unique genes encoding PPP enzymes (*tal, rpiA, rpe*; (**Appendix Fig S1**) on bacterial growth in different planktonic conditions and in J774.1 macrophages (**Fig 3A, B**). For this, we used three transposon insertion mutants obtained from the *F. novicida* Tn insertion library (Gallagher *et al*, 2007). The three mutants were designated Δ*rpe*, Δ*rpiA* and Δ*tal*, for simplification.

**Figure 3.**
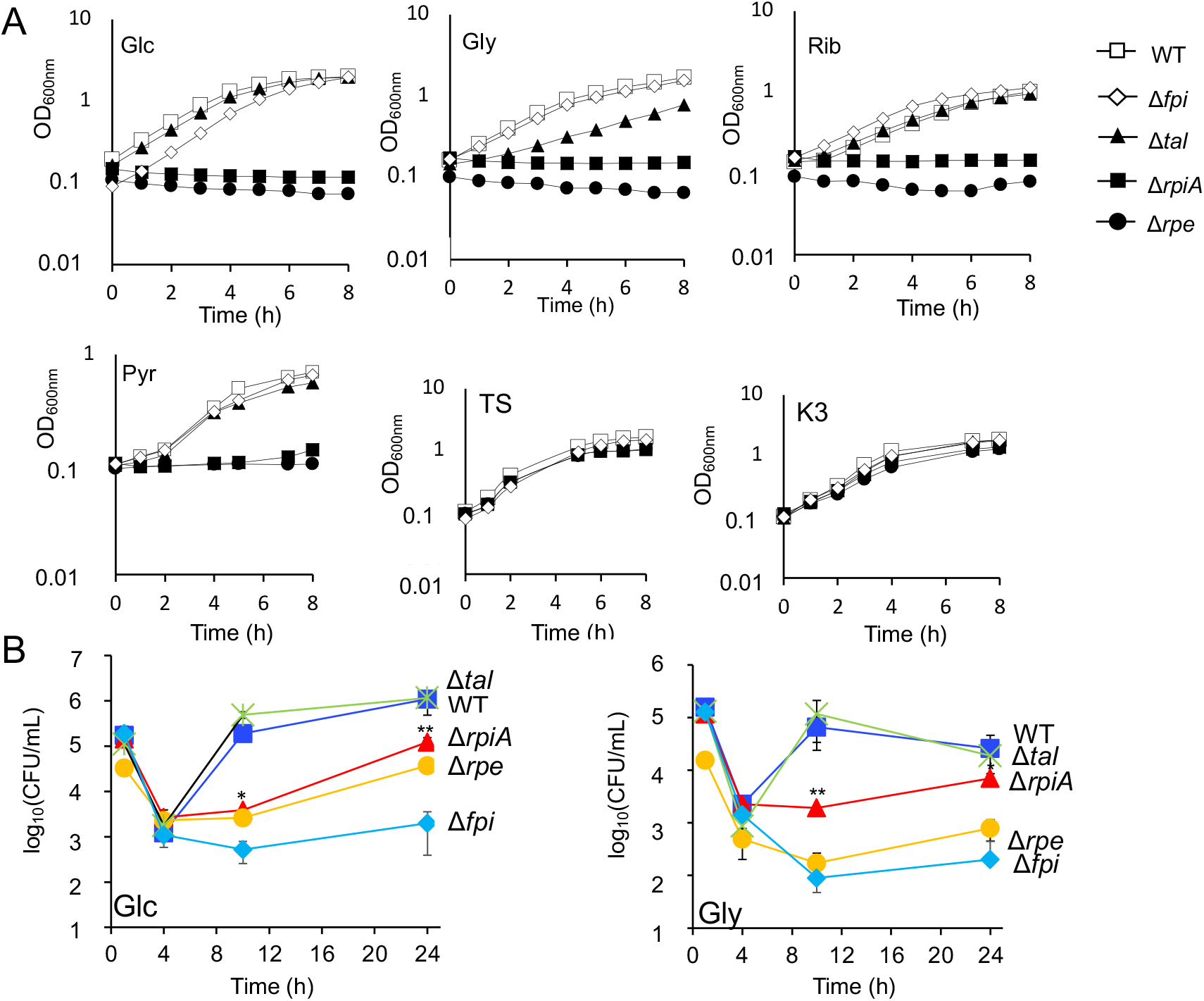
Growth of *ΔrpiA, Δrpe and Δtal* PPP mutants in liquid culture. (**A**) Bacterial growth was monitored in chemically defined medium (CDM), supplemented with various carbon sources (Glc, glucose; Gly, glycerol; Rib, ribose; Pyr, pyruvate), each at a final concentration of 25 mM; as well as in tryptic soy broth supplemented with cysteine (0.1% w/v) and glucose (0.4% w/v) (TS); or Shaedler K3 medium (K3). Stationary-phase bacterial cultures of wild-type *F. novicida* (WT), *ΔrpiA, Δrpe and Δtal* mutants were diluted to a final OD_600nm_ of 0.1, in 20 mL broth. Every hour, the OD_600nm_ of the culture was measured, during a 9 h-period. (**B**) Kinetics of intracellular multiplication of the three PPP mutants Δ*rpe*, Δ*rpiA* and Δ*tal*. Intracellular multiplication of the three mutants was monitored in J774.1 over a 24 h-period, in DMEM supplemented with either Glucose (Glc, left panel) or Glycerol (Gly, right panel), and compared to that in the wild-type *F. novicida* (WT). A Δ*fpi* mutant strain was used as a negative control.

Two mutants, Δ*rpe* and Δ*rpiA*, failed to grow in any of CDMs tested confirming that the PPP is essential for growth in synthetic media. In contrast, the Δ*tal* mutant only showed a modest growth defect in CDM supplemented with carbon sources likely to integrate the glycolytic/gluconeogenic axis (glycerol and fructose). All three mutants showed wild-type growth in rich media (TS and K3), indicating that they do not correspond to essential genes. The intracellular growth properties of Δ*rpe*, Δ*rpiA* and Δ*tal* mutants were therefore next monitored in J774.1 macrophages. Since we previously showed that the intracellular behavior of *F. novicida* mutants could vary depending on the carbon source supplemented in the DMEM medium (Brissac et al 2015; Ziveri et al. 2017), bacterial multiplication was monitored in macrophages grown in the presence of either glucose or glycerol (**Fig 3B**). In DMEM-glucose, the Δ*rpiA* and Δ*rpe* mutants multiplied only poorly during the first 10 hours post-infection but a restart of intracellular multiplication was recorded at 24 h, for both mutants. However, the colony forming units (CFUs) recorded were still approximately 8- to 30-fold lower than that of WT. Of note, in DMEM-glycerol, the Δ*rpiA* mutant showed better multiplication than the Δ*rpe* mutant. In contrast, the Δ*talA* mutant showed WT multiplication in both media, and at all time-points tested, suggesting the conservation of a transaldolase activity in this mutant. Of note, since each of the intermediate metabolites of the PPP can theoretically be synthesized in the absence of Tal, this enzyme might be dispensable under most physiological conditions.

### Dynamics of macrophage infection of the PPP mutants

We then used imaging approaches to better characterize, at the cellular level, the impact of the four mutations in the PPP.

We first checked the subcellular localization of the four PPP mutants, using GFP-labeled bacteria, and monitored their colocalization with the late phagosomal marker LAMP-1 by confocal immunofluorescence microscopy, at 10 h and 24 h (**Fig 4A**). As expected, the Δ*fpi* negative control showed a high colocalization with LAMP-1 at both time points (75% and 80% at 10 h and 24 h, respectively), confirming that it remained trapped into phagosomes. In contrast, only 5.3% colocalization was recorded with wild-type *F. novicida*, at both time points. At 10 h, the Δ*tktA*, Δ*rpiA*, Δ*rpe* and Δ*tal* mutants showed 22.3%, 18%, 10.9% and 7.6%, of colocalization with LAMP-1 respectively, indicating that, for all four mutants, the majority of the bacteria had escaped the phagosomal compartment. Colocalization of the four mutants remained very low after 24 h (7.4%, 10.7%, 12.1% and 4.8%, respectively). (**Fig 4B**). These results suggest that the observed intracellular growth defect of the four PPP mutants is mainly a consequence of their inability to cope with the environment of the cytosolic compartment of infected macrophages.

**Figure 4.**
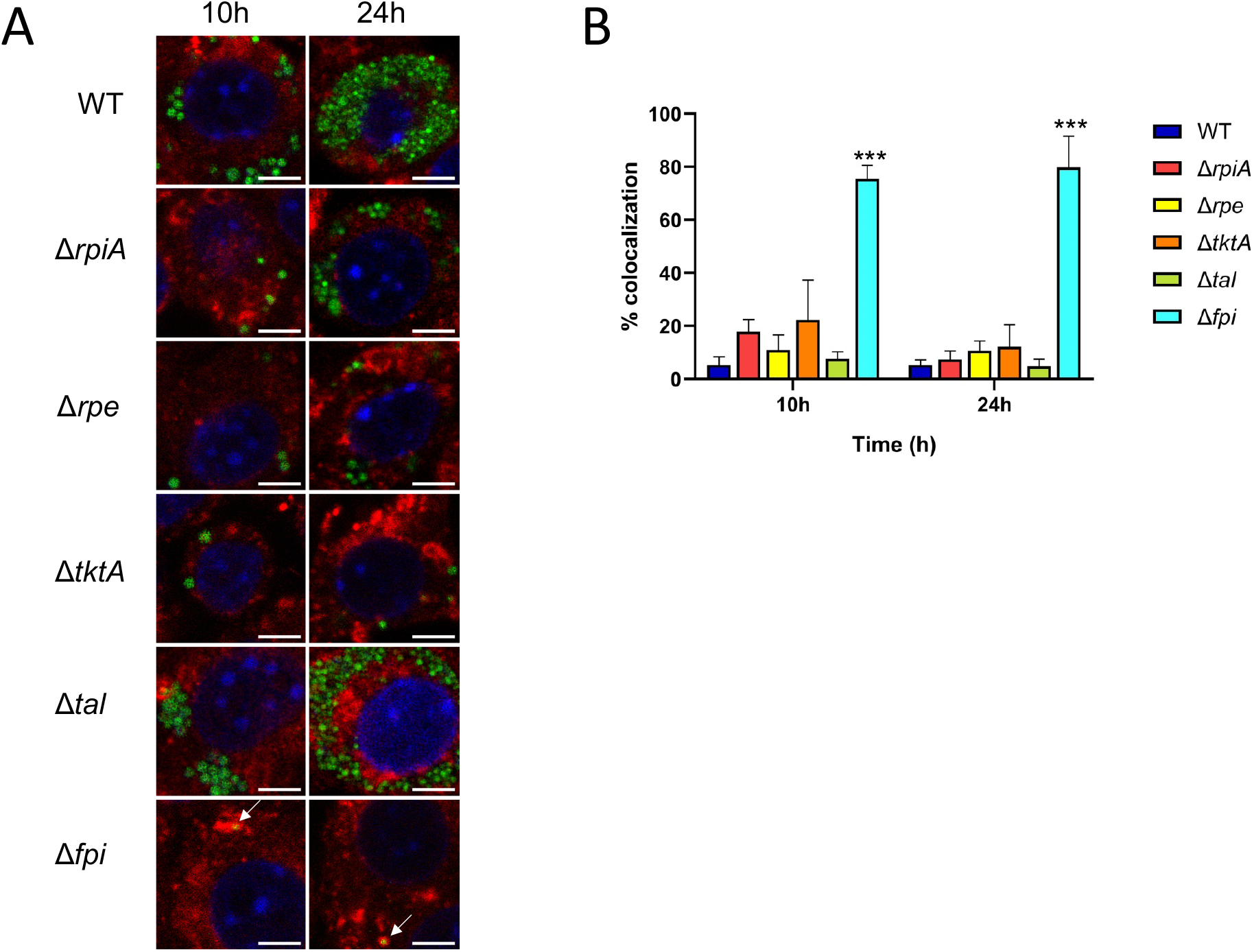
Subcellular localization of the PPP mutants. Glucose-grown J774.1 were infected for 1 h with wild-type *F. novicida* (WT), Δ*rpiA*, Δ*rpe*, Δ*tktA*, Δ*tal*, or Δ*FPI* strains and their co-localization with the phagosomal marker LAMP-1 was observed by confocal microscopy 10 and 24 h, after beginning of the experiment. (**A**) *Scale bars* at the *bottom right* of each panel correspond to 5 μM. J774.1 were stained for *F. tularensis* (*green*), LAMP-1 (*red*), and host DNA (*blue*, DAPI stained). (**B**) Quantification of bacteria/phagosome colocalization in glucose-grown J774.1 macrophages. Mean and SD of triplicate wells. ∗∗∗p<0,0001 (determined by ANOVA one-way test). The white arrows point to Δ*fpi* bacteria entrapped in a phagosomal compartment (orange).

We next wished to visualize, at the single cell level, the intracellular multiplication in the four PPP mutants. For this, we used time-lapse video microscopy, using a fully automated microscope (Incucyte® 531 S3, Essen BioScience), J774–1 macrophages with red nuclei (here designated J774.1_red_ and GFP-expressing bacteria (see Materials and Methods). J774.1_red_ cells were infected by GFP-expressing bacteria at an MOI of 100 and infection was followed over a 48h-period, in 96-well plates (**Fig 5, Appendix Fig S3** and **Movie E1 to E5**). We followed the total number of J774.1_red_ cells over time to evaluate the impact of bacterial infection on cell viability (see Materials and Methods). This assay showed that cells were able to continuously multiply at least during the 40 first hours and were not affected by bacterial infection (**Fig 5A**). Intracellular bacterial multiplication was followed by monitoring the total green area intensity (reflecting multiplication of the GFP-positive bacteria) over time (**Fig 5B**) and was translated on graph line by using the Incucyte S3 software. Multiplication of the Δ*tktA*, Δ*rpe* and Δ*rpiA* mutants was affected to variable extents and the Δ*tktA* mutant was again the most severely affected whereas the Δ*tal* mutant showed wild-type multiplication. The total number of GFP-positive J774.1_red_ cells was next followed by monitoring the total number of cells with red nuclei associated with at least one detectable signal of green intensity (see Materials and Methods). The total number of J774.1_red_ cells infected with wild-type bacteria increased during the first 10 h of infection, then reached a plateau until 40h and increased again between 40 and 48 h (**Fig 5C** and **Fig S4**). This increase was faster and continuous with the Δ*tal* mutant than with the wild-type strain. In contrast with the three PPP mutants, the total number of J774.1_red_ cells infected with GFP-expressing bacteria progressively decreased with time, suggesting a continuous elimination of these mutant bacteria similar to that of the Δ*fpi* negative control strain. At selected time points (10 h, 24 h, and 48 h), the amount of GFP signal detected was accurately quantified (GFP area, see Materials and Methods). This analysis revealed that at 10h, more than 85% of the GFP-positive cells contained few bacteria as indicated by a GFP measured area of less than 200 pixels for all tested strains (estimation range 1 to 10 bacteria per cell; **Fig 5D**). However, in 1-2% of the cells infected with Δ*rpiA* and Δ*rpe* mutants some bacterial multiplication had occurred since the GFP measured area were in the 200-400 pixels range. As expected, wild-type bacteria exhibited a greater and heterogenous intracellular multiplication capacity with GFP measured area also detected in the 200-400, 400-600 and up to the 600-800 pixels range. Remarkably, the Δ*tal* mutant also showed a heterogeneous and even broader bacterial intracellular multiplication capacity than the wild-type strain.

**Figure 5.**
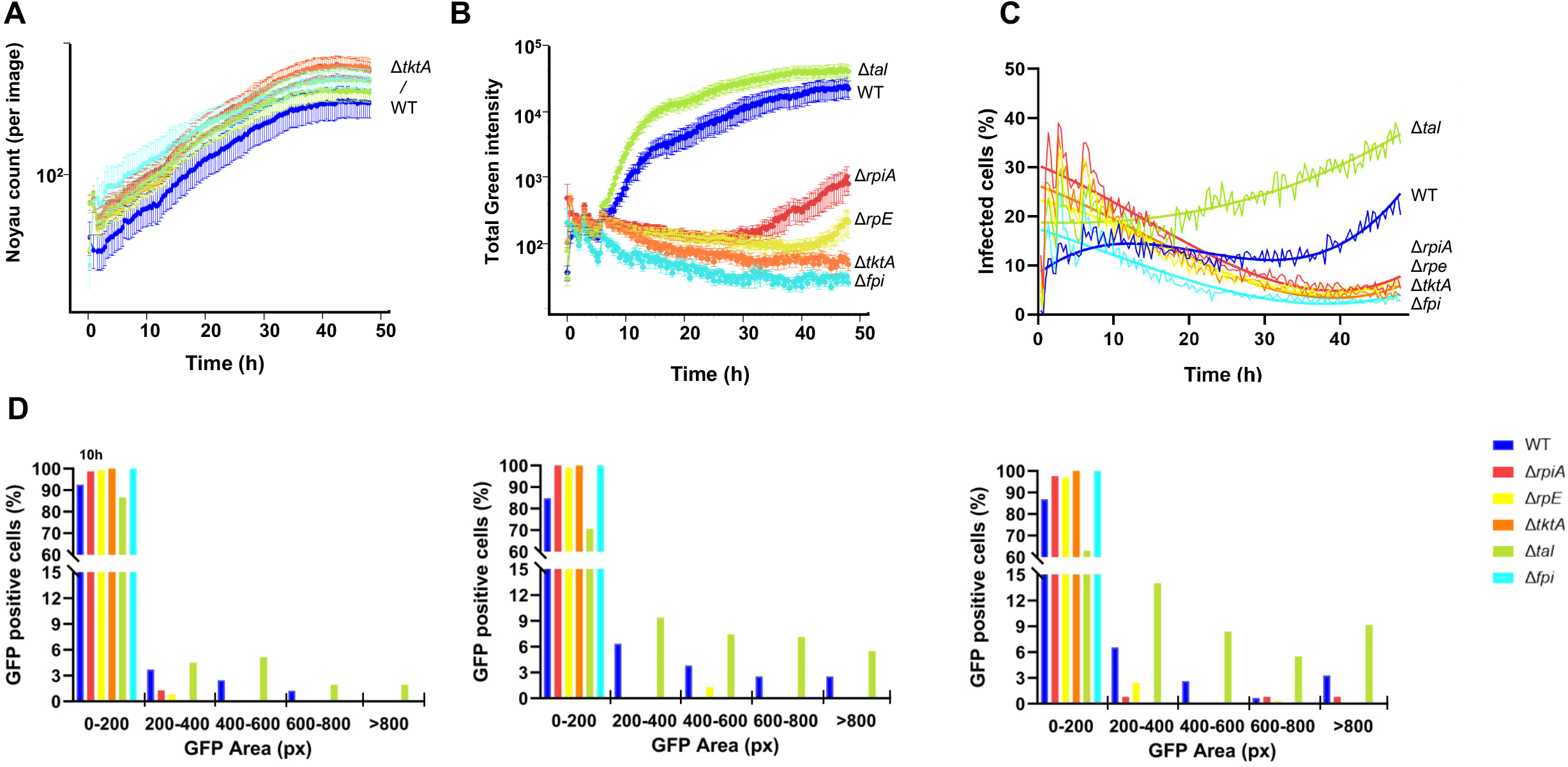
Time lapse video microscopy analyses of the PPP mutants. Kinetics of intracellular multiplication of the PPP mutants generated by the Incucyte S3 software (***Incucyte*** Live-Cell Analysis System, Sartorius). Strains were transformed by GFP-carrying plasmid and intracellular multiplication of the four mutants was recorded by video-microscopy in J774.1_red_ over a 48 h-period, in DMEM supplemented with Glucose and compared to that in the wild-type F. novicida (WT). A Δfpi mutant strain was used as a negative control. (A) The total number of J774.1_red_ cells was monitored over time was determined by the Incucyte S3. (B) Bacterial multiplication (GFP-expressing bacteria) was followed by monitoring the total green area intensity over time. Multiplication was translated on graph line by using Incucyte S3, Sartorius. (C) The total number of GFP-positive cells was followed by monitoring the total number cells with red nuclei associated with at least one detectable signal of green intensity (*i*.*e*. infected with GFP-expressing bacteria). (D) At selected time points (10 h, 24 h, and 48 h), the number of GFP-positive cells were decomposed into 5 categories depending on the intensity of the GFP signal detected.

At 24 h, approximately 30% and at 48 h, 40% of the cells infected with the Δ*tal* mutant showed a GFP area >200, as compared to only 15% with the wild-type strain. Of note, at 24h, cells infected with Δ*rpiA* and Δ*rpe* mutants were still identified in the 200-400 pixels range (1 −2% of the cells) and for the Δ*rpiA* mutant approximately 2% of the cells were in the >600 pixels range, revealing an active multiplication of this mutant in a limited subset of cells. Overall these analyses, which fully supported CFUs counts, revealed that Δ*tktA*, Δ*rpe* and Δ*rpiA* mutants had impaired intracellular multiplication until 30 h after infection. The Δ*rpe* and Δ*rpiA* mutants then resumed growth in a limited subset of cells, reaching up to 30% that of WT. A very modest intracellular multiplication of the Δ*tktA* mutant could also be visualized between 40 h and 48 h (**Fig 5 and Movie E1 to E5**). The Δ*tal* mutant showed wild-type or even improved intracellular multiplication-dissemination, in all the conditions tested.

### Virulence assay in the adult fly

Finally, the impact of PPP mutations was evaluated *in vivo*. We used the adult fly model, a simple animal model which has been previously used to evaluate attenuation of virulence of banks of *Francisella* mutants (Asare *et al*, 2010).

Adult male flies were infected with wild-type *F. novicida* (WT), or Δ*tktA*, Δ*rpiA*, Δ*rpe*, Δ*tal*, and Δ*fpi* isogenic mutants and monitored fly survival over a 10 day-period (see Materials and Methods). 100% of the flies infected with either the WT or the Δ*tal* strain died within 8 days whereas 100% of the flies infected with the Δ*fpi* mutant strain survived. The Δ*tktA*, Δ*rpiA* and Δ*rpe* mutants were only slightly less virulent than the WT and killed 90% of the flies after 10 days. Among them, the Δ*tktA* mutant killed significantly more slowly than the WT strain (**Appendix Fig S5**). Hence, in spite of showing impaired intracellular multiplication, the Δ*tktA*, Δ*rpiA* and Δ*rpe* mutants retained most of their virulence properties in this model.

Having established the impact of PPP genes inactivation on bacterial virulence, we then decided to undertake a thorough proteo-metabolomic analysis of the PPP mutants, in order to: i) deepen our understanding of the consequences of impaired PPP activity on the pathophysiology of *F. novicida*; and ii) to identify links between PPP and other metabolic pathway possibly contributing to intracellular niche adaptation.

### Invalidation of PPP key enzymes leads to common variations of the global proteomic profile

We first performed a whole-cell comparative nanoLC-MS/MS proteomic analysis of WT, and the three mutant strains that showed impaired intracellular growth, *ΔtktA, Δrpe and ΔrpiA*. As a control we also performed the whole cell comparative analysis of WT and of the Δ*tal* mutant that did not show any intracellular growth defect (**Fig 6)**. In all cases, invalidation of the gene, and therefore absence of the protein, induced a strong modulation of the global amount of protein compared to the WT.

**Figure 6.**
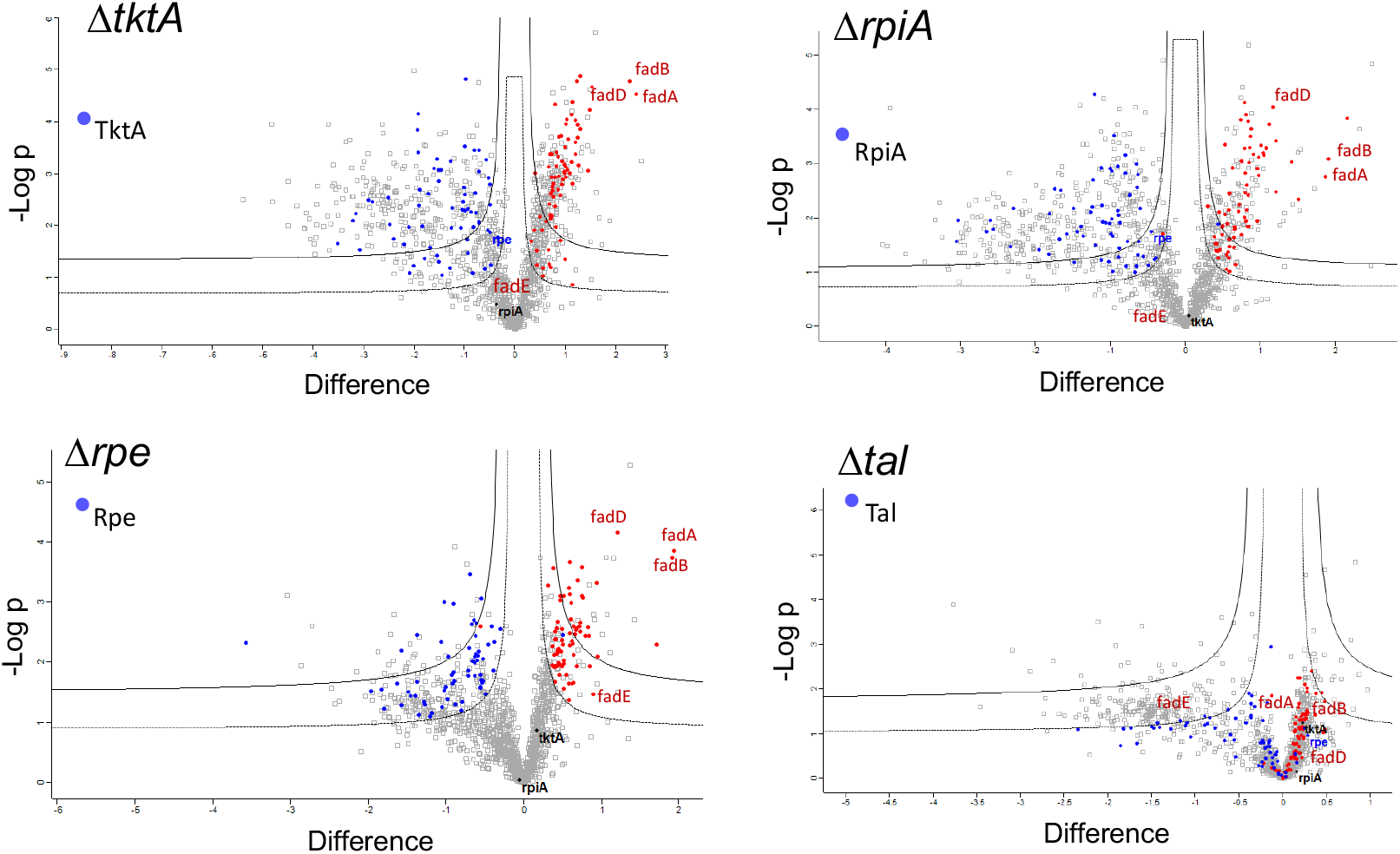
Differential proteomes of WT and Δ*tktA, ΔrpE and ΔrpiA* mutants. The proteomes of the four PPP mutants Δ*tktA*, Δ*rpiA*, Δ*rpe* and Δ*tal*, was compared to that of WT *F. novicida*. Volcano plot representing the statistical comparison of the protein LFQ intensities of each mutant versus WT. Inner volcano was established using S0 = 0.1, FDR = 0.05 and the outer volcano using S0 =0.1, FDR =0.01. The abscissa reports the fold change in logarithmic scale (difference), the ordinate the –log(pvalue). Proteins undergoing the same modulation in Δ*tktA*, Δ*rpe* and Δ*rpiA* mutants but not in Δ*talA* are highlighted in color (blue and red for decreased increased in the mutant, respectively).

Globally, 1 502 proteins were identified across samples. Of those, we retained 1 381 proteins confidently quantified in at least one condition, covering 80% of the *Francisella* proteome (out of 1 722 proteins predicted to be encoded by the *F. novicida* U112 genome). We performed a t-test (FDR<0.05) showing that a large number of proteins was impacted by the deletions: 835 proteins by Δ*tktA*, 690 proteins by Δ*rpiA*, 372 by Δ*rpe* and 237 by Δ*tal* (**Appendix Table S2**). As expected, for each mutant, the proteins deleted from the genome were found as absent in the corresponding mutant. Of note, in all three mutants Δ*tktA*, Δ*rpiA* and Δ*rpe* mutants, expression of the Tal protein was found increased.

By comparing the proteins modulated in each mutant, we observed that there was a common set of 145 proteins that was modulated in the four mutants Δ*tktA*, Δ*rpiA*, Δ*rpe* and Δ*tal*. However, an additional set of 137 proteins was modulated in Δ*tktA*, Δ*rpiA*, Δ*rpe* mutants but not in the Δ*tal* mutant which might be involved in their impaired intracellular behavior (**Fig 6** and **Appendix Table S2**).

The vast majority of these proteins was following the same modulation pattern, with 73 upregulated and 64 downregulated in the three mutant strains. Of note, the amplitude of protein modulation, in terms of fold changes, was stronger in Δ*rpiA* and Δ*tktA*, than in Δ*rpe*.

Of note, most of the proteins encoded by the fatty acid degradation locus *fad*, were up-regulated in the three mutants but not in the Δ*tal* control strain **(Appendix Fig S6)**. These results regarding the corresponding *fad* genes were confirmed by qRT-PCR in the Δ*tktA* mutant indicating that the impact of *tktA* gene inactivation on Fad proteins expression occurred already at the transcriptional level. Indeed, a ca. 3-fold increased expression of most of the genes of the *fad* operon was recorded in exponentially-grown bacteria in Shaedler K3 medium (OD_600_ 0.5), in the mutant compared to WT in late exponential phase (OD_600_ 1), this increase reached up to 6-fold that of WT. These data strongly suggest a link between PPP and fatty acid metabolism.

### Metabolomic analyses of the PPP mutants

Inactivation of the *tktA* gene resulted in important changes in global metabolome. Fifty metabolites are presented (**Fig7)**. Forty metabolites levels changed significantly (23 down and 17 up). An increase of metabolites belonging to different chemical classes was recorded, notably amino acids, nucleotides and sugars. The glucose oxidation intermediates were significantly affected with marked accumulation of glucose, the glycolytic intermediate F-1,6P, as well as the PPP metabolite Ribose-P. It also yielded the accumulation of dihydroxyacetone phosphate (DHAP), a metabolite tightly associated to the glycolytic/gluconeogenic pathways. Indeed, DHAP is a breakdown product of fructose 1,6-bisphosphate (F-1,6P) by the enzyme fructose biphosphate aldolase (FBA) and can be converted to GA-3P by the enzyme triose phosphate isomerase. Of note, in *E. coli*, mutations in *transketolases* have been shown to lead to an accumulation of DHAP, a precursor of the highly toxic compound methylglyoxal (Girgis *et al*, 2012). On the contrary *tktA* inactivation was accompanied by decreased concentrations of the PPP intermediate sedoheptulose-7P (S-7P) as well as of the glycolytic end product pyruvate.

In line with what was observed with Δ*tktA, rpiA* inactivation resulted in the accumulation of the glycolytic intermediate G-6P, and of Ribose-P. Like the Δ*tktA* mutant, this mutant also yielded the accumulation of DHAP. However, in sharp contrast to Δ*tktA*, Δ*rpiA* inactivation resulted in increased levels of S-7P. *rpiA* inactivation was also accompanied by decreased production of the glycolytic intermediate GA-3P. *rpe* inactivation resulted in similar effect as *rpiA* with accumulation of S-7P and glucose. In addition, *rpe* inactivation resulted in decreased pyruvate. Overall Δ*rpiA* and Δ*rpe* inactivation manifested a similar metabolic phenotype that was distinct from that of the Δ*tktA* mutant.

In addition, all the mutants shared a relative decrease of long chain fatty acids (LCFAs), such as: i) palmitic and stearic acids (16 and 18 carbon, saturated LCFAs, respectively), in both Δ*tktA* and Δ*rpiA* mutants; ii) palmitoleic acid (16 carbon, monounsaturated LCFA), in Δ*rpe* and Δ*rpiA* mutants; or iii) oleic acid (18 carbon, monounsaturated FA), in Δ*tktA* and Δ*rpe* mutants. Hence, metabolic profiling of mutants inactivated in the PPP showed common traits with alterations in the relative amounts of similar glycolytic intermediates and fatty acids.

These data are also consistent with proteomic and transcriptomic analysis showing an upregulation of the whole *fad* operon and further confirm the link between the PPP and fatty acid metabolism.

### Integration of the Proteometabolomic data

We anticipated that *Francisella* would reprogram its metabolism, in response to the loss of PPP enzymatic activity. Regularized Canonical Correlation Analysis (rCCA) was performed and metabolite–protein networks were identified, at a highly stringent correlation threshold of 0.95 (**Fig 8** and **Appendix Fig S7**). Networks were performed for each pair mutant/WT, separately (Δ*tktA*/WT; Δ*rpiA/*WT and Δ*rpe*/WT). Given the linear correlation of variables used in rCCA, a correlation in the same direction was called positive and corresponded either to a decreased level of the metabolite associated with a decreased expression of the protein or to an increased level of the metabolite associated with an increased expression of the protein. Conversely, when the levels of the metabolite and the protein vary in the opposite direction, it was called a negative correlation.

**Figure 7.**
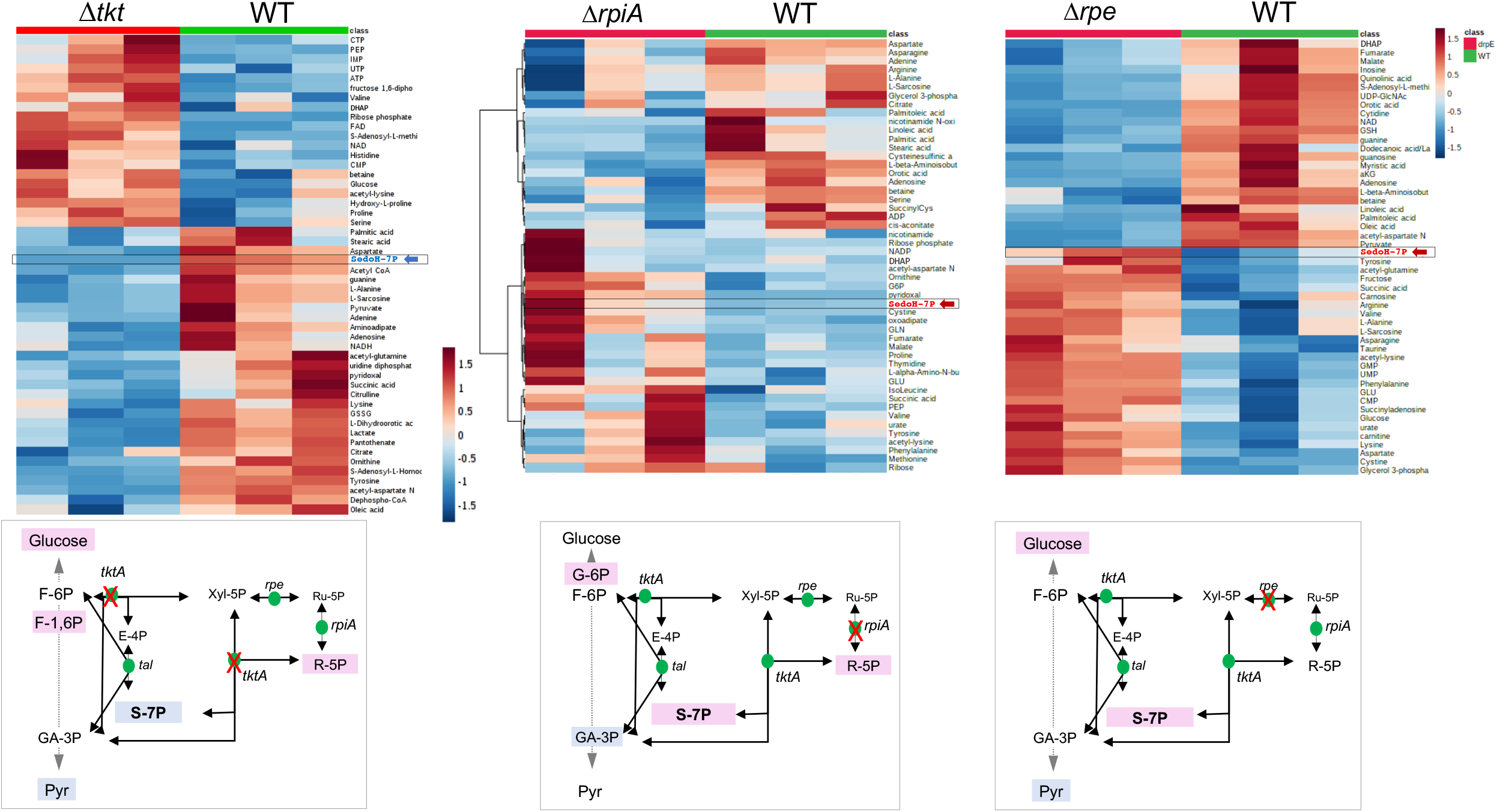
Comparison of metabolite profiles of WT and Δ*tktA, Δrpe and ΔrpiA* mutants. The isolates were cultured to the stationary phase in TSB supplemented with cysteine (0,1% w/v)) and glucose (0,4% w/v). Heatmap visualization and hierarchical clustering analysis of the metabolite profiling in each mutant compared to WT *F. novicida*. Upper part, heatmaps showing the top 50 (Δ*tktA, Δrpe*) or top 30 (*ΔrpiA*) most changing compounds. Three biological replicates, performed for each sample, are presented. Rows: metabolites; columns: samples; color key indicates metabolite relative concentration value (blue: lowest; red: highest). The arrows, to the right of each heatmap, pinpoint the metabolites related to the PPP or glycolytic pathways. Lower part, the position of the metabolites related to the PPP or glycolytic pathways is shown on a schematic depiction of the pathways.

**Figure 8.**
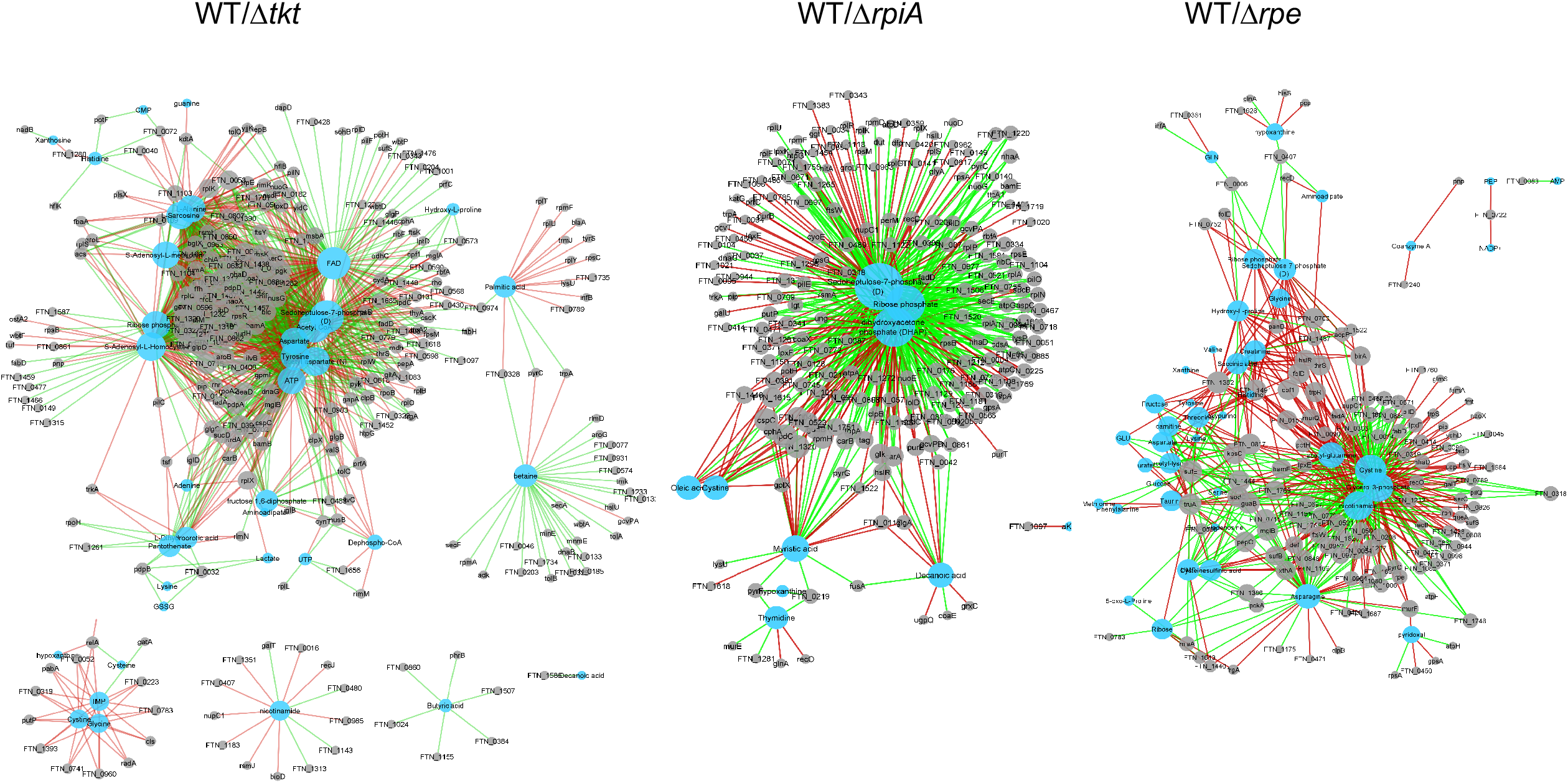
Correlation network derived from rCCA between metabolites and proteins from *Francisella*. (**A**) WT/Δ*tktA*; (**B**) WT/Δ*rpiA*; (**C**): WT/Δ*rpe*.

For each network, three major metabolic hubs were identified, each linking one metabolite to multiple proteins. For the Δ*tktA/*WT network we achieved a cross-validation score (cv-score) = 0.9879 with λ1 = 0.1 and λ2 = 1.788889. The biggest hub (150 correlated proteins) was Sedopheptulose-7P (S-7P), with 80 proteins positively correlated and 70 proteins negatively correlated. The second and third largest hubs were Flavin Adenine Dinucleotide (FAD) (144 correlated proteins) and acetyl-CoA (133 correlated proteins), respectively. Of note, all three metabolites can be correlated to carbohydrate catabolic processes: S-7P as a central metabolite of the PPP; FAD as an electron carrier participating to both catabolic and anabolic reactions; and acetyl-CoA as the end product of glycolysis. For the Δ*rpiA/*WT network the cv-score was 0.9017 with λ1 = 0.0001 and λ2 = 0.9. The biggest hub (191 corelated proteins) was also Sedopheptulose-7P (S-7P), with 63 proteins positively correlated and 128 proteins negatively correlated. The second and third largest hubs were DHAP (134 corelated proteins) and the PPP metabolite ribose-P (117 corelated proteins), respectively. Of note, whereas in Δ*tktA* mutant, expression of S-7P was decreased compared to WT, in the Δ*rpiA* mutant S-7P was increased compared to WT. In this case also, all three metabolic hubs can be correlated to carbohydrate catabolic processes: S-7P and Ribose-P to the PPP; and DHAP to the glycolytic/gluconeogenic pathways. Finally, for the Δ*rpe/*WT network the cv-score was 0.9502 with λ1 = 0.0001157895 and λ2 = 0.0001894737. The biggest hub (104 corelated proteins) was cystine, with 60 proteins positively correlated and 44 proteins negatively correlated. The second and third largest hubs were the glycolytic intermediate GA-3P (93 corelated proteins) and Nicotinamide Adenine Dinucleotide (NAD, an electron carrier like FAD, 87 corelated proteins), respectively. In contrast to the two other networks, the biggest hub (cystine) has no obvious link to carbohydrate metabolism or the PPP. However, the two other hubs, *eg*. GA-3P (a metabolite at the crossroad of glycerol metabolism, glycolysis and the PPP) and NAD, participating to both catabolic and anabolic reactions (including glycolysis and the tricarboxylic acid cycle), can be associated to these processes. The cysteine related hub was associated with 104 proteins, including approximately 80% proteins belonging to different predicted functional categories (*i*.*e*. amino acid metabolism and transport, replication and repair, translation, lipid metabolism and cell wall membrane /envelope biogenesis) and 20% proteins of unknown function. As expected, Rpe was found negatively correlated with cystine (since cystine increased and the Rpe protein decreased in the Δ*rpe*/WT network).

In both Δ*tktA*/WT and Δ*rpiA*/WT networks, S-7P was associated with a large number of proteins (150 to 191) displaying a broad spectrum of predicted biological activities (**Fig 8** and **Appendix Fig S7**). Notably in both cases, the largest category of proteins (except the proteins of unknown functions) was associated to the translation machinery, including the ribosomal proteins L11, L15, L17, L23, L24 of the 50S large subunit and S4, S13, S18, S30 of the 30S small subunit, for the network Δ*tktA*/WT(positively correlated since both were decreased); and L6, L11, L15, L19, L21, L24, L30 of the 50S subunit and S1, S2, S5, S17, S13 of the 30S subunit, for the network Δ*rpiA/*WT(negatively correlated since S-7P increased and the ribosomal proteins decreased). As expected, TktA was found positively correlated with S-7P (since both are decreased in the Δ*tktA*/WT network) and RpiA was found negatively correlated with S-7P (since S-7P in increased and the RpiA protein decreased in the Δ*rpiA*/WT network)

Overall, integration of our proteometabolic data highlighted links between PPP and other metabolic pathways by revealing major and/or new hubs such as S-7P and cystine. We hypothesize that such important proteometabolic hubs could greatly impact *Francisella* pathogenicity.

## Discussion

We show here that the PPP constitutes a crucial metabolic hub for intracellular multiplication of *Francisella*. Our data suggest that the PPP is instrumental during the early phase of the intracellular life cycle.

### Temporal contribution of the PPP to intracellular multiplication

Bacteria, which possess both oxidative and non-oxidative branches of the PPP, are able to coordinate glycolysis/gluconeogenesis with the PPP to control the production of either NADPH or Ribose-5 Phosphate (R5-P), according to the environmental conditions. For example, *E. coli* can reroute its glycolytic flux into the oxidative branch of the PPP to achieve the immediate replenishment of NADPH for glutathione reduction upon oxidative stress (Stincone *et al*, 2015). In contrast, bacteria such as *Francisella* and *Legionella*, which both lack the oxidative branch, use other pathways to produce NADPH.

In *Francisella* as well as in *L. pneumophila* (Häuslein *et al*, 2017) and *C. burnetii*, the gene *tktA*, encoding transketolase, and genes encoding enzymes of the glycolysis/gluconeogenesis pathways (*gapA, pgK, pyk*), are organized in an operon (**Fig 1** and **Appendix Fig S2**). In *L. pneumophila*, the regulation of this operon has been shown to be under the control of the RNA binding protein CsrA (Sahr *et al*, 2017). Several other pathogenic bacterial species also have *tktA, gapA* and *pgk* genes in the same cluster and with the same organization but lack the *pyk* and *fba* genes. This is the case of *Bordetella pertussis*, and *Brucella melitensis*. In most *Burkholderia* species (including *B. multivorans, B. pseudomallei, B. thailandensis*, …), the *tktA* and *gapA* genes are adjacent and in the same orientation, suggesting that they belong to the same transcriptional unit, whereas the *pgk, pyk*, and *fba* gene cluster is located in a distinct region of the chromosome (**Appendix Fig S2)**.

We found that expression of the *tktA* and *gapA* genes was comparable in all the planktonic conditions tested. Indeed, *gapA* was only 1.5- to 1.7-fold more transcribed than *tktA* in CDM-glucose or CDM-glycerol in early exponential phase whereas, in macrophages, the expression of *gapA* was approximately 5-fold higer higher than that of *tktA*, suggesting that the glycolytic/gluconeogenic metabolic axis may be permanently required during bacterial intracellular multiplication and at a higher level of activity than the PPP.

We and others have shown that gluconeogenesis, but not glycolysis, was used as the main pathway for host-derived amino acids utilization as carbon and nitrogen sources. (Brissac *et al*, 2015; Ziveri *et al*, 2017b; Radlinski *et al*, 2018). In addition, Kawula and co-workers have recently demonstrated that host cell lipolysis was also required during *Francisella* intracellular replication, suggesting that host-derived triglyceride stores (*i*.*e*. esters derived from glycerol and three fatty acids) may represent a primary source of glycerol for the bacterium in the cytosolic compartment.

We found here that inactivation of three genes of the PPP (*tktA, rpe* and *rpiA*) provoked an initial intracellular growth arrest up to 10 h after infection, whereas at later time points, bacterial multiplication started to resume, especially with the Δ*rpe* and Δ*rpiA* mutants. These data led us to propose a model (**Fig 9)** in which intracellular bacteria necessitate both PPP and glycolysis during their initial phase of multiplication. However, at later time points, when gluconeogenesis becomes the major route of host-derived nutrients utilization, the PPP might be no longer essential.

**Figure 9.**
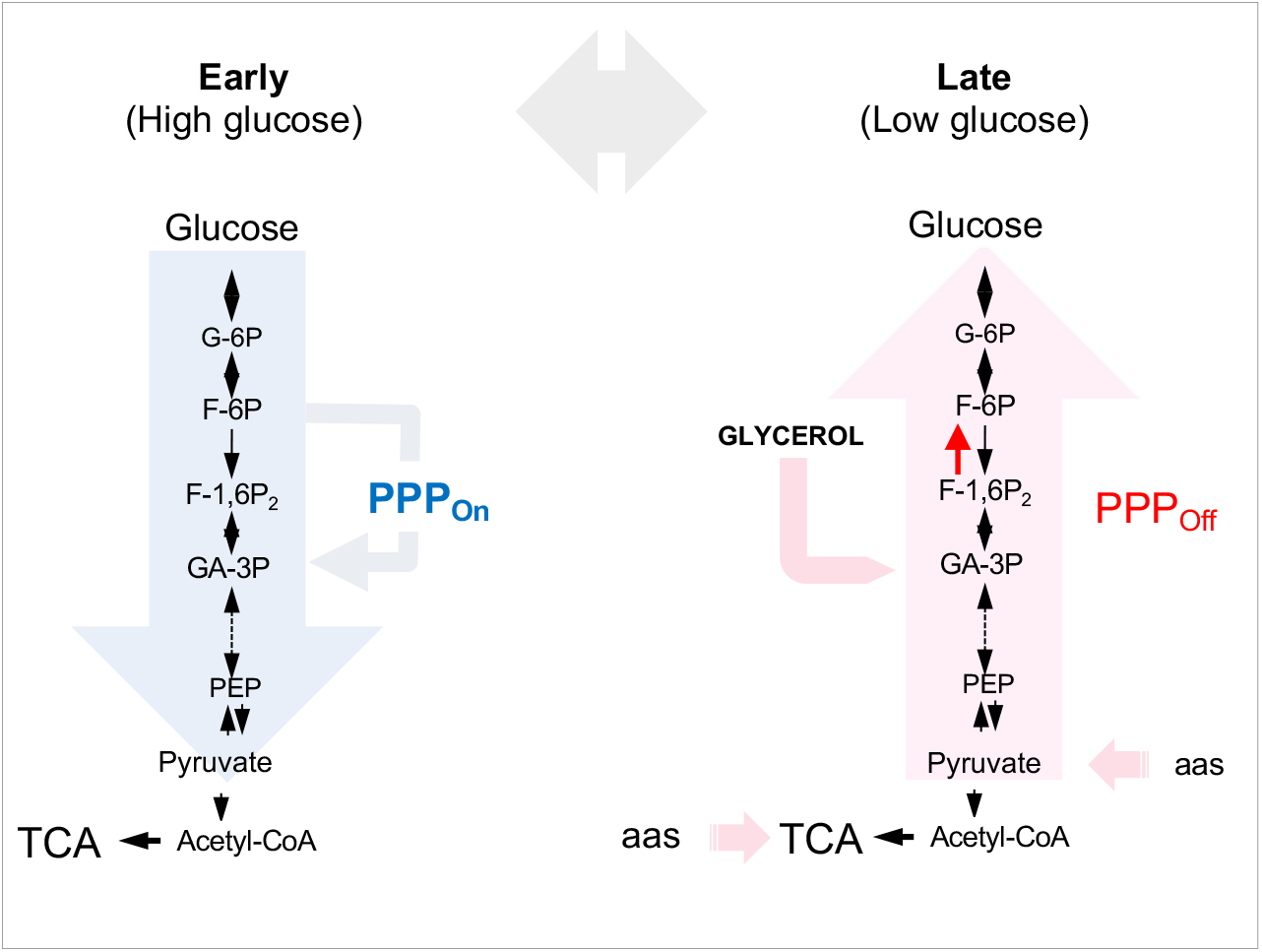
Model of PPP and glycolysis/gluconeogenesis interplay during macrophage infection. Left panel (early), during the first 24 h of the intracellular life cycle, when glucose is still available in replete condition, Glycolysis (blue arrow) and the PPP are “On”. Right panel (late), at later time points (24 − 48 h), when glucose becomes limiting, gluconeogenesis prevails (pink arrow) and the PPP turns “Off”.

### Cross-talk between PPP and other metabolic pathways

Our earlier fluxomic analyses have shown that a glycolytic flux through the PPP existed (Brissac *et al*, 2015). In contrast, in the presence of the gluconeogenic substrate pyruvate, compounds of the PPP were not detected, suggesting that the gluconeogenic flux from pyruvate did not involve the participation of the PPP. Carbon flux analyses performed by Eisenreich and co-workers (Chen *et al*, 2017) confirmed that exogenous glucose was the most efficient carbon substrate for *F. novicida* when growing in complex medium and suggested a possible glycolytic turnover via the PPP and/or gluconeogenesis.

Both Tal and TktA enzymes act as a bridge between glycolysis and the PPP by sharing intermediate metabolites with glycolysis (F-6P and GA-3P) (Takayama *et al*, 1997). Somewhat unexpectedly, loss of tal-encoded transaldolase activity had no deleterious impact on *Francisella* intracellular fate and even improved its intracellular multiplication. Interestingly, Tal has been shown to play a key role in PPP in some bacteria but not others (Nakahigashi *et al*, 2009; Koendjbiharie *et al*, 2020). Hence, either *Francisella* possesses a functional paralog lacking significant amino acid sequence similarity with Tal, or another enzyme known to play a different role, may also be capable of compensating, under certain metabolic conditions, for the absence of transaldolase.

The fact that the three other PPP mutants (Δ*tktA*, Δ*rpe* and Δ*rpiA*) were still able to survive and replicate after 24 h might be explained by the rescue provided by the pool of host-derived intermediates. Indeed, it has been shown in *Chlamydia* that nucleotide transport proteins (NTTs) catalyzed the import of nucleotides from the eukaryotic host into the bacterial cell and rendered de novo synthesis of these compounds dispensable (Knab *et al*, 2011; Gehre *et al*, 2016). Nucleotide utilization was also recently described in parasites (Major *et al*, 2019) Microsporidia, a group of strict intracellular eukaryotic parasites which cannot make nucleotides, also use host-derivied nucleotides. For this, they are equipped with multiple paralogs of the major facilitator superfamily (MFS) transporters to exploit the host nucleotide pool. *Francisella* also possesses multiple transporters (Alkhuder *et al*, 2010), including phagosomal transporter (Pht) family members (Fonseca *et al*, 2014; Sauer *et al*, 2005; Ramond *et al*, 2015; Marohn *et al*, 2012). Of note, in *L. pneumophila*, two of these transporters, PhtC and PhtD, were proposed to contribute to protect *the bacterium* from dTMP starvation during its intracellular life cycle.

Several intracellular pathogens including, *Listeria, Legionella, Coxiella*, and *Chlamydia*, have been shown to develop a bi-modal way of using nutrients coined “bipartite metabolism”(Best & Abu Kwaik, 2019), contributing to the temporal and spatial intracellular niche adaptation. Whereas *L. pneumophila* and *C. burnetii* species use host amino acids, and especially serine, as primary sources of carbon and energy, *L. monocytogenes* and *C. trachomatis* use host glycerol or malate, respectively, to feed their TCA cycle and *L. monocytogenes* uses glucose and glucose-6P primarily for biosynthesis of cell wall components and nucleotides.

### Novel metabolic networks

Our laboratory has recently shown that, in the Gram-positive pathogen *Staphylococcus aureus, tktA* inactivation led to dysregulation of the whole cell metabolism (Tan *et al*, 2019). As in *Francisella*, an accumulation of R-5P and a decrease in S-7P amount were recorded in the Δ*tktA* mutant compared to wild-type *S. aureus*. Interestingly, a very recent study by Sachla and Helmann showed that the toxic metabolite 4-P-erythronate (4PE) was accidentally produced from the promiscuous reaction of GapA with erythrose-4-phosphate of the PPP (Sachla & Helmann, 2019), illustrating the multiple, and sometimes unexpected, connections between glycolysis and the PPP.

Integrative Omic approaches are important to have a global view of the changes triggered upon inactivation of individual genes (Jean Beltran *et al*, 2017). Network biology is based on the principle that biological processes are not chiefly controlled by individual molecules or by discrete, unconnected linear (Charitou *et al*, 2016). Here, we applied rCCA to explore correlation structures between metabolites and proteins, to explore the links between the PPP and other metabolic pathways, including glucolysis/gluconeogenesis, in *Francisella*, as well as to discovered novel metabolic crosstalks and connecting hubs.

This allowed us to identify several putative novel connections of the PPP with other pathways and notably with the Fatty acid degradation pathway (Fad) and with cystine metabolism. Our whole cell proteomic approach revealed that the Fad pathway was upregulated in the mutants Δ*tktA*, Δ*rpiA* and Δ*rpe*. qRT-PCR analyses of the *fad* locus further confirmed its upregulation in the Δ*tktA* mutant compared to WT *Francisella* **(Appendix Fig S6)**. In *E. coli*, the *fad* operon is under the negative control of the FadR repressor whose promoter binding activity has been shown to be altered upon binding of long chain unsaturated fatty acid. FadR has no ortholog in *Francisella* genomes. Hence, it is likely that the regulation of *fad* genes in *Francisella* is different from that in *E. coli*.

Metabolomic analyses also revealed a decrease in the amounts of several long chain and middle chain fatty acids in the metabolomes of the three mutants, compatible with their increased degradation. It is tempting to suggest that this increased fatty acid degradation might be a mean for the mutant bacteria to increase the pool of acetyl-CoA (the end-product of the pathway) in order to rapidly fuel the TCA and compensate for the lack of the PPP.

Of particular interest, rCCA integration of proteomic and metabolic data of the Δ*rpe*/WT pair identified cystine as a major hub. *Francisella* is auxotrophic for cysteine and thus absolutely relies on external sources of cysteine for growth. We and others have previously shown that (Alkhuder *et al*, 2009; Ramsey *et al*, 2020) glutathione (γ-Glutamate-cysteine-glycine or GSH) constituted a major host-derived source of cysteine for intracellular *Francisella*. Cystine, which results from the dimerization of cysteine, is also available and can be used as sources of sulfur. This indispensable chemical element is required for the activity of many enzymes and is involved in ion and redox metabolic pathways (Burguière *et al*, 2004). Of note, three proteins involved in sulfur metabolism are correlated to cystine in the Δ*rpe*/WT pair: SufS, SufB (FTN_0851, Fe-S cluster assembly protein), and SufE. In *E. coli*, SufE together with the SufBCD complex, has been shown to enhance SufS cysteine desulfurase activity, as part of a sulfur transfer pathway for Fe-S cluster assembly (Outten *et al*, 2003). The connection to the PPP remains to be elucidated since none of the enzymes of glycolysis or the PPP rely on Fe-S clusters.

The integration of proteomics and metabolomics data open the way to the discovery of novel metabolic crosstalks contributing and/or controlling the intracellular life cycle of *Francisella* and should apply to other intracellular bacterial pathogens, offering thus new perspectives on host-pathogen interplay.

## Materials and Methods

### Ethics Statement

All Materials and Methods involving animals were conducted in accordance with guidelines established by the French and European regulations for the care and use of laboratory animals (Decree 87–848, 2001–464, 2001–486 and 2001–131 and European Directive 2010/63/UE) and approved by the INSERM Ethics Committee (Authorization Number: 75-906).

### Strains and culture conditions

All strains used in this study are derivative from *F. tularensis* subsp. *novicida* U112 (here designated *F. novicida*). Strains were grown at 37°C on PolyViteX agar plates (BioMerieux), in Tryptic Soy Broth (TSB) or Chemically Defined Medium (CDM). The Chemically Defined Medium used for *F. novicida* corresponds to standard CDM (Chamberlain, 1965) without threonine and valine (Gesbert *et al*, 2014). For growth condition determination, bacterial strains were inoculated in the appropriate medium at an initial OD_600nm_ of 0.1 from an overnight culture in TSB. The *tktA* mutant was constructed by allelic replacement (see below). The *rpiA, rpe, tal*, transposon insertion mutants of *F. novicida* and the Δ*fpi* mutant were kindly provided by Anders Sjöstedt (Umea University, Sweden).

### Construction of a Δ*tktA* deletion mutant

We inactivated the gene *tktA* in *F. novicida* (*FTN_1333*) by allelic replacement resulting in the deletion of the entire gene. Briefly, we generated by overlap PCR a recombinant PCR product containing the upstream region of the gene *tktA* (*tktA* -UP), a kanamycin resistance cassette (*nptII* gene fused with *pGro* promoter) and the downstream region of the gene *tktA* (*tktA* -DN). Primers *tktA* upstream FW (p1) and *tktA* upstream (splK7) RV (p2) amplified the 700 bp region upstream of position + 1 of the *tktA* coding sequence (*tktA*-UP), primers *pGro* FW (p3) and *nptII* RV (p4) amplified the 1091 bp kanamycin resistance cassette (*nptII* gene fused with *pGro* promoter); and primers *tktA* downstream (splK7) FW (p5) and *tktA* downstream RV (p6) amplified the 491 bp region downstream of the position +1991 of the *tktA* gene coding sequence (*tktA*-DN). PCR reactions were realized using Phusion High-Fidelity DNA Polymerase (ThermoScientific) and PCR products were purified using NucleoSpin® Gel and PCR Clean-up kit (Macherey-Nagel). The overlap PCR product was purified from agarose gel and was directly used to transform wild type *F. novicida* by chemical transformation (Gesbert et al., 2015). Recombinant bacteria were isolated by spreading onto Chocolate agar plates containing kanamycin (10μg mL^-1^). The mutant strain was checked for loss of the wild type gene by PCR product direct sequencing (GATC-biotech) using appropriate primers.

### Functional complementation

The plasmid pKK-*tktA*_*cp*_, used for complementation of the *F. novicida* Δ*tktA* (Δ*tktA*), is described below. Primers pX-FW *tktA* and pY-RV *tktA* amplified the 236 bp region immediately upstream of *tktA* start codon and the complete *tktA* gene from U112. The PCR product was purified and SmaI/PstI restricted in presence of FastAP Thermosensitive Alkaline Phosphatase (ThermoScientific) to avoid self-ligation, and cloned into pKK214 vector after SmaI/PstI double restriction and transformed in *E. coli* TOP10. Recombinant plasmid pKK-*tktA*_*cp*_ was purified and directly used for electroporation in *F. novicida* Δ*tktA* (Gesbert *et al*, 2015). Recombinant colonies were selected on PolyViteX agar plates containing tetracycline (5μg mL^-1^) and kanamycin (10μg mL^-1^).

### Proteomic analyses

#### Protein digestion

FASP (Filter-aided sample preparation) procedure for protein digestion was performed as previously described, using 30 kDa MWCO centrifugal filter units (Microcon, Millipore, Cat No MRCF0R030).

Briefly, sodium dodecyl sulfate (SDS, 2% final) was added to 30 μg of each lysate to increase solubility of the proteins, in a final volume of 120 μL. Proteins were reduced with 0.1M dithiotreitol (DTT) for 30 min at 60 °C, then applied to the filters, mixed with 200 μL of 8M urea, 100mM Tris-HCl pH 8.8 (UA buffer), and finally centrifuged for 15 min at 15,000 x g. In order to remove detergents and DTT, the filters were washed twice with 200 μl of UA buffer. Alkylation was carried out by incubation for 20 min in the dark with 50mM iodoacetamide. Filters were then washed twice with 100 μl of UA buffer (15,000 x g for 15 min), followed by two washes with 100 μl of ABC buffer (15,000 x g for 10 min), to remove urea. All centrifugation steps were performed at room temperature. Finally, trypsin was added in 1:30 ratio and digestion were achieved by overnight incubation at 37 °C.

#### NanoLC-MS/MS protein identification and quantification

Samples were vacuum dried, and resuspended in 30 μL of 10% acetonitrile, 0.1% trifluoroacetic acid for LC-MS/MS. For each run, 1 μL was injected in a nanoRSLC-QExactive PLUS (RSLC Ultimate 3000, ThermoScientific, Waltham, MA, USA). Peptides were separated on a 50 cm reversed-phase liquid chromatographic column (Pepmap C18, Thermo Scienfitic). Chromatography solvents were (A) 0.1% formic acid in water, and (B) 80% acetonitrile, 0.08% formic acid. Peptides were eluted from the column with the following gradient of 120 min. Two blanks were run between triplicates to prevent sample carryover. Peptides eluting from the column were analyzed by data dependent MS/MS, using top-10 acquisition method. Briefly, the instrument settings were as follows: resolution was set to 70,000 for MS scans and 17,500 for the data dependent MS/MS scans in order to increase speed. The MS AGC target was set to 3Å∼106 counts, while MS/MS AGC target was set to 1Å∼105. The MS scan range was from 400 to 2000m/z. MS and MS/MS scans were recorded in profile mode. Dynamic exclusion was set to 30 s duration. Three replicates of each sample were analyzed by nanoLC-MS/MS.

#### Data processing following nanoLC-MS/MS acquisition

The MS files were processed with the MaxQuant software version 1.5.8.30 and searched with Andromeda search engine against the *Uniprot F. novicida* database (release 2016, 1 722 entries). To search parent mass and fragment ions, we set a mass deviation of 3 and 20 ppm respectively. The minimum peptide length was set to 7 amino acids and strict specificity for trypsin cleavage was required, allowing up to two missed cleavage sites. Carbamidomethylation (Cys) was set as fixed modification, whereas oxidation (Met) and N-term acetylation were set as variable modifications. The false discovery rates at the protein and peptide levels were set to 1%. Scores were calculated in MaxQuant as described previously. The reverse and common contaminants hits were removed from MaxQuant output. Proteins were quantified according to the MaxQuant label-free algorithm using LFQ intensities; protein quantification was obtained using at least 1 peptide per protein.

Statistical and bioinformatic analysis, including heatmaps, profile plots, and clustering, were performed with Perseus software (version 1.5.5.31) freely available at www.perseus-framework.org73. For statistical comparison, we set two groups, WT and Δ*tktA*, each containing four biological replicates. Each sample was run in technical triplicates as well. We then filtered the data to keep only proteins with at least 3 valid values out 4 in at least one group. Next, the data were imputed to fill missing data points by creating a Gaussian distribution of random numbers with a SD of 33% relative to the SD of the measured values and 2.5 SD downshift of the mean to simulate the distribution of low signal values. We performed an T test, FDR<0.001, S0 = 1.

Hierarchical clustering of proteins that survived the test was performed in Perseus on logarithmic scaled LFQ intensities after z-score normalization of the data, using Euclidean distances.

### Metabolomic analyses

Metabolites profiling of *F. novicida* isolates was performed by liquid chromatography–mass spectrometry (LC-MS) as described (Mackay *et al*, 2015). Two independent experiments with three biological replicates were performed for each isolate. Briefly, *F. novicida* isolates were grown in CFSM with addition of thymidine in the same conditions as for the proteomic analyses. Metabolic activity was blocked by immersion in liquid nitrogen for 10 sec.

Metabolites were extracted using a solvent mixture of Methanol/ACN/H_2_O (50:30:20) at −20 °C. Samples were vortexed for 5 min at 4°C, and then centrifuged at 16,000 g for 15 minutes at 4°C. The supernatants were collected and analyzed by LC-MS using SeQuant ZIC-pHilic column (Millipore) for the liquid chromatography separation. The aqueous mobile-phase solvent was 20 mM ammonium carbonate plus 0.1% ammonium hydroxide solution and the organic mobile phase was acetonitrile. The metabolites were separated over a linear gradient from 80% organic to 80% aqueous phase for 15 min. The column temperature was 48 °C and the flow rate was 200 μl/min. The metabolites were detected across a mass range of 75–1,000 m/z using the Q-Exactive Plus mass spectrometer (Thermo) at a resolution of 35,000 (at 200 m/z) with electrospray ionization and polarity switching mode. Lock masses were used to ensure mass accuracy below 5 ppm. The peak areas of different metabolites were determined using TraceFinder software (Thermo) using the exact mass of the singly charged ion and known retention time on the HPLC column.

Statistical analyses were performed using MetaboAnalyst 4.0 software. The algorithm for heatmap clustering was based on the Pearson distance measure for similarity and the Ward linkage method for biotype clustering. Metabolites with similar abundance patterns were positioned closer together.

### Network analysis

Aiming the metabolomics and proteomics data integration, we used the regularized canonical correlation analysis (rCCA) to build the correlation networks. The rCCA is a modification of the classical canonical correlation analysis (CCA), which is a multivariate statistical method used to assess correlations between two multivariate datasets (Gonzalez *et al*, 2008). Three biological replicates of each dataset were used (proteomics and metabolomics) and, in order to reduce the discrepancy of the scales, we combined some transformations and scaling methods, like generalized logarithm transformation (Durbin *et al*, 2002), pareto scaling (van den Berg *et al*, 2006) or z-score. CCA aims to maximize the correlation between linear combinations of variables (canonical variates) in two datasets (Hong *et al*, 2013). The rCCA analysis used is part of the mixOmics package V5.2 (Lê Cao *et al*, 2009) for R (http://www.R-project.org). Regularization parameters (λ_1_ and λ_2_) were estimated using the *tune*.*rcc()* function to evaluate the cross-validation score (*cv-score*) for each point in the network, achieving the values for λ_1_ and λ_2_ that offered the highest *cv-score*. To create the canonical correlations and the canonical variates between the datasets we use the *rcc()* function and the *network()* to produce the networks. We set the correlation threshold to ≥ 0.95, this value was chosen to obtain biologically interpretable networks that were neither too sparse nor too dense (González *et al*, 2012). After, the networks were exported to Cytoscape network visualization software V3.8 (Shannon *et al*, 2003), using the organic layout with diameter of the node relative to the number of undirected edges. The intensity of edge colors represent the correlation values ranging from dark green (negative correlation) to dark red (positive correlation). Since hubs are network structures that could play a key role in biological networks (Montastier *et al*, 2015; Hong *et al*, 2013), we focus in the three biggest hubs of each network and the direct correlations of proteins related to the *fad* operon.

The number of positive and negative correlations and the complete description of each network are reported in the **Appendix Table S3** and the cytoscape (.cys) files of each network are provided in **Appendix Table S4**.

### Cell cultures and cell infection experiments

J774A.1 (ATCC® TIB-67™) cells were propagated in Dulbecco’s Modified Eagle’s Medium (DMEM, PAA), containing 10% fetal bovine serum (FBS, PAA) unless otherwise stated. For CFU counting, the day before infection, approximately 2.10^5^ eukaryotic cells per well were seeded in 12-wells cell tissue plates and bacterial strains were grown overnight in 5 mL of TSB at 37°C.

Infections were realized at a multiplicity of infection (MOI) of 100 and incubated for 1 h at 37°C in culture medium. After 3 washes with cellular culture medium, plates were incubated for 4, 10 and 24 h in fresh medium supplemented with gentamycin (10 µg mL^-1^). At each kinetic point, cells were washed 3 times with culture medium and lysed by addition of 1 mL of distilled water for 10 min at 4°C. Viable bacteria titers were determined by spreading preparations on chocolate plates. Each experiment was conducted at least twice in triplicates.

### Fly infection

For each test, 20 adult male *drosophila* were infected by being pricked with a glass needle dipped into bacterial colonies of the wild type or mutant *F. novicida* strains. The flies were incubated at 29°C and transferred to fresh food daily. Living animals were counted once a day for 10 days, and the results were recorded as a percentage of the number of living animals relative to the number of flies recorded one day post-pricking. Flies pricked by a clean needle were considered as control of the experiment. Two independent replicates were carried out. The Gehan–Breslow–Wilcoxon test was used to compare the killing efficiency of the different mutants to that of the wild-type *F. novicida* strain.

### Transcriptional analysis

#### Isolation of total RNA and reverse transcription

For transcriptional analyses of bacteria grown in CDM supplemented either with glucose or glycerol, cultures were centrifuged for 2 min in a microcentrifuge at room temperature and the pellet was quickly resuspended in Trizol solution (Invitrogen, Carlsbad, CA, USA). For transcriptional analyses of bacteria in infected cells, J774.1 macropahges grown in standard DMEM-glucose medium were infected with wild-type *F. novicida* (WT) strain for 24 h. Cells were then collected by scratching, centrifuged at maximum speed in a microcentrifuge at room temperature and the pellet was quickly resuspended in Trizol solution. Samples were either processed immediately or frozen in liquid nitrogen and stored at −80 °C. Samples were treated with chloroform and the aqueous phase was used in the Monarch® RNA Cleanup Kit (New England Biolabs Inc).

#### Quantitative real-time RT-PCR

WT *F. novicida* and mutant strains were grown overnight at 37 °C. Then, samples were harvested and RNA was isolated and reverse trancripted in cDNA following protocol manufacturer (LunaScript™RTSuperMix Kit, New England Biolabs Inc). The 20 μL reaction consisted in 4 μL of RT Mix completed with 16 μL of water with 100 ng of RNA. qPCR was performed according to manufacturer’s protocol on Applied Biosystems-ABI PRISM 7700 instrument (Applied Biosystems). We used Luna® Universal qPCR Master Mix (New England Biolabs, Inc), following protocol manufacturer, by adding 1 μL from RT mix. Transcript levels were analyzed using a 7900HT Fast Real-Time PCR System (Applied Biosystems) according to the standard settings of the system software. The thermal cycling conditions were: 50°C for 2 min, 95°C for 10 min, followed by 40 cycles of 95°C for 15 s and 60°C for 1 min. We used the “Relative Standard Curve Method” to analyze qRT-PCR data. The amounts of each transcript were normalized to helicase rates (*FTN_1594*).

### *In silico* analyses

For synteny prediction, we used the Kyoto Encyclopedia of Genes and Genomes (KEGG) Sequence Similarity Database (SSDB) to search conserved gene clusters containing homologs of *FTN_1333* using default parameters (gap size 0, threshold 100).

The SSDB database (Release 94.0, last accessed on 14th April 2020) contains SSEARCH computation results (based on the Smith-Waterman similarity score) for all pairwise genome comparisons of the KEGG database (10.1093/nar/gkw1092).

The Microbial Genomic context Viewer MGcV allowing the coloring of the genes by Clusters of orthologous gene (COG) category retrieved from NCBI RefSeq was used for visualization (10.1186/1471-2164-14-209).

### Confocal experiments

J774.1 macrophages were infected (MOI of 1,000) with wild-type *F. novicida* U112 (WT), the pentose phosphate pathway mutants (Δ*tktA*, Δ*rpe*, Δ*rpiA*, Δ*tal*), or an isogenic strain deleted for the “Francisella Pathogenicity Island” (ΔFPI) in standard DMEM (DMEM-glucose) for 30 min at 37°C. All the bacterial strain carried a GFP-carried-plasmid. Cells were then washed three times with PBS and maintained in fresh DMEM supplemented with gentamycin (10μg mL^-1^) until the end of the experiment. Three kinetic points (1 h, 10 h and 24h) were sampled. For each point cells were washed with 1X PBS, fixed 15 min with 4% Paraformaldheyde, and incubated 10 min in 50 mM NH_4_Cl in 1X PBS to quench free aldehydes. Cells were then blocked and permeabilized with PBS containing 0.1% saponin and 1% bovin serum albumin for 10 min at room temperature. Cells were then incubated for 30 min with anti-LAMP-1 rabbit polyclonal antibody (1/100 final dilution, ABCAM) After washing, cells were incubated for 30 min with Alexa546 conjugated donkey anti rabbit secondary antibodies (1/400e, AbCam). After washing, DAPI was added (1/10 000e) for 1 min and glass coverslips were mounted in Mowiol (Cityfluor Ltd.). Cells were examined using an X63 oil-immersion objective on a Leica SP8 gSTED confocal microscope. Co-localization tests were quantified by using Image J software; and mean numbers were calculated on more than 100 cells for each condition. Confocal microscopy analyses were performed at the Cell Imaging Facility (Faculté de Médecine Necker Enfants-Malades).

### Time lapse video microscopy

#### Constructions of GFP-expressing strains

Plasmid pKK-pGro-GFP (Ziveri *et al*, 2019) was introduced by chemical transformation into wild-type *F. novicida* (designated WT-GFP) and in the Δ*tktA*, Δ*rpiA*, Δ*rpe* and Δ*tal* mutants, to generate strains that constitutively expressing GFP.

#### Time-lapse microscopy

The three GFP-expressing strains were imaged using the IncuCyte™ technology. J774.1_red_ cells were grown to confluence in 96-well cell tissue plates and were infected with GFP-expressing wild-type and mutant bacteria, at a MOI of 100. Plates were then incubated for 1 h at 37 °C in culture medium. After 3 washes with cellular culture medium, plates were incubated at 5% CO2 and 37 °C for 48 h in fresh medium with 5% of SVF supplemented with gentamycin (10 g ml^-1^). Bacterial multiplication was monitored in the fully automated microscope Incucyte® S3 (Essen BioScience). Images were taken every 20 min with the 20X objective. Green and red fluorescence images were obtained every 20 min with an acquisition time of 400 milliseconds (ms) and 200 ms respectively. Time-lapse videos (from which images were extracted) were generated by using Incucyte® S3 and imageJ softwares.

#### Data analyses

Raw images of the phase representing the cells as well as green fluorescence representing the GFP bacteria were extracted for each time point. First, we used the opensource program IlastiK v1.3.3post3 to train the machine to recognize the cells from the phase images. The same method was used to define and recognize the bacteria via the green fluorescence images. For each category (cells or bacteria), a mask was extracted from the analysis. A segmentation by size filter was carried out, using the Fiji program, on the “cells” mask in order to exclude the cellular debris. Then, the cells and bacterial masks were superimposed. This procedure allowed to: i) measure the total number of cells, at each time point, and ii) for each cell, at each time point, the presence of a GFP signal (bacteria) and the GFP area were measured (**Appendix Fig S4**). The assay was performed on 8 wells for each condition.

### Statistics

*In vitro* experiments were at least repeated twice and in triplicates. Data were analyzed using GraphPad Prism software. Tests are specified in each legend. In figures, all the results correspond to mean ± SEM.

## Data Availability

The mass spectrometry proteomics data have been deposited to the ProteomeXchange Consortium via the PRIDE partner repository with the dataset identifier PXD023550. The authors declare that all other data supporting the findings of this study are available within the paper and its Supplementary Information files.

## Acknowledgements

We thank Dr A. Sjostedt for providing the *Francisella* strains: wild-type *F. novicida* U112, its Δ*fpi* mutant derivative, and the Δ*tal*, Δ*rpe* and Δ*rpiA* transposon insertion mutants (*FTN_0781, FTN_1221, FTN_1185*, respectively). These studies were supported by INSERM, CNRS and Université de Paris. Jason Ziveri and Héloise Rytter funded by a fellowship from the “Délégation Générale à l’Armement”. The funders had no role in study design, data collection and analysis, decision to publish, or preparation of the manuscript.

## Appendix PDF

Supplementary Information includes 7 Supplementary figures (**Appendix Fig S1 to S7**) and 4 supplementary tables (**Appendix Table S1 to S4**).

**Appendix Figure S1**. Schematic depiction of the major steps of the PPP and glycolytic/glucoenoegenic pathways.

**Appendix Figure S2**. (**A**) *tktA* gene cluster synteny analysis within representative *Francisella* genomes. Comparative context map of *FTN_1333* homologs obtained via KEGG Sequence Similarity Database (SSDB) search is visualized using MGcV. Genes belonging to the same Clusters of orthologous gene (COG) groups are depicted in the same color. COG color key: light green: Carbohydrate transport and metabolism; dark green: Lipid transport and metabolism; red: Translation, ribosomal structure and biogenesis; grey: Hypothetical proteins. (**B**) *tktA* gene cluster synteny analysis within representative genomes of selected plant and human pathogens. Comparative context map of *FTN_1333* homologs obtained via KEGG Sequence Similarity Database (SSDB) search is visualized using MGcV. Genes belonging to the same Clusters of orthologous gene (COG) groups are depicted in the same color. COG color key: light green: Carbohydrate transport and metabolism; dark green: Lipid transport and metabolism; red: Translation, ribosomal structure and biogenesis; grey: Hypothetical proteins; white: unknown function; yellow: Energy production and conversion; brown: Inorganic ion transport and metabolism; orange: transcription; blue: Nucleotide transport and metabolism.

**Appendix Figure S3**. Screenshots of time lapse video microscopy.

Pictures of GFP-expressing wild-type *F. novicida* (WT), Δ*tal*, Δ*rpiA*, Δ*rpe*Δ, *tktA* or Δ*fpi* strains, were taken one hour after infection of J774.1_red_ cells and gentamycin washing (0 h), 10 h and 24 h after intracellular survival. Bar = 50 µm.

**Appendix Figure S4**. Bioinformatic analysis of time-lapse images.

(**A**) The open source program IlastiK v1.3.3post3 recognize the cells from the phase images and the bacteria from the green fluorescence images, respectively. (**B**) A segmentation by size filter was carried out, using the Fiji program, on the “cells” mask in order to exclude the cellular debris. Then, the cells and bacterial masks were superimposed.

**Appendix Figure S5**. Virulence in the adult fly model.

Survival curves of adult male flies infected by WT strain, Δ*tkt*, Δ*rpiA*, Δ*rpe*, Δ*tal*, Δfpi mutant at 29 °C. Results were analyzed by Gehan–Breslow–Wilcoxon test (∗∗P < 0.01).

**Appendix Figure S6**. (**A**) Schematic organization of the *fad* operon. (**B**) qRT-PCR analysis.

(**C**) Schematic representation of the metabolic pathway leading to acetyl-CoA production. (**D**) Proteometabolic analysis of *fad*-encoded proteins and their relationship to metabolite changes in Δ*tkt*, Δ*rpiA* and Δ*rpe* mutants.

**Appendix Figure S7**. Families of proteins identified with the major metabolic hub of the Δ*tkt*,

Δ*rpiA*, Δ*rpe* mutants.

**Appendix Table S1**. List of 135 differentially expressed in the 3 mutants Δ*tktA*, Δ*rpiA* and

Δ*rpe*, compared to wild-type *F. novicida* (WT).

**Appendix Table S2**. This rCCA summary lists the number of proteins associated with the 3 major hubs of each pair Mutant/WT.

**Appendix Table S3**. List of FTN_1333 homologs with proteins predicted to be in a gene cluster according to KEGG SSDB.

**Appendix Table S4**. Cytoscape files. Each file has the main network, the three biggest hubs, and the *fad* locus.

**Appendix Movies E1 to E5**. Wild-type *F. novicida* (E1), Δ*rpiA* (E2), Δ*rpe* (E3), Δ*tktA* (E4), Δ*tal* (E5).

